# Reduction of collinear inhibition in observers with central vision loss using anodal transcranial direct current stimulation: A case series

**DOI:** 10.1101/2020.11.09.375600

**Authors:** Rajkumar Nallour Raveendran, Amy Chow, Katelyn Tsang, Arijit Chakraborty, Benjamin Thompson

## Abstract

People with central vision loss (CVL) due to macular degeneration are forced to rely on their residual peripheral vision and often develop a preferred retinal locus (PRL), a region of intact peripheral retina that is used for fixation. At the PRL, visual processing is impaired due to crowding (cluttering of visual objects). The problem of crowding still persists when images are magnified to account for the lower resolution of peripheral vision. We assessed whether anodal transcranial direct stimulation (a-tDCS), a neuro-modulation technique that alters cortical inhibition, would reduce collinear inhibition (an early component of crowding) when applied to the visual cortex in patients with CVL. Our results showed that applying a-tDCS to the visual cortex for 20mins reduced crowding in three patients with CVL and that the effect was sustained for up to 30mins. Sham stimulation delivered in a separate session had no effect. These initial observations mandate further research into the use of a-tDCS to enhance cortical processing of residual retinal input in patients with CVL.

## Introduction

Macular degeneration can be caused by eye diseases that occur early in life (juvenile macular degeneration; JMD) or in older adulthood (wet or dry age-related macular degeneration; AMD). AMD will affect about 3 million individuals in the United States by 2020 [1,2]. JMD is less common (1-2 in 10,000 individuals)[3] but the impact is greater due to a life-long loss of vision. Current treatments for macular degenerative diseases can stabilize or slow disease progression but generally cannot provide a ‘cure’. As a result, a significant number of individuals with macular degeneration suffer from a loss of central vision that forces them to rely on para-central peripheral vision. Most patients with macular degeneration (84% of 1339 eyes)[4] develop a preferred retinal locus (PRL): a specific para-central retinal region that is used for visual tasks in place of the fovea. However, the para-central retina has lower spatial resolution (poorer visual acuity) than the fovea. More importantly, the para-central retina is more susceptible to crowding (an inability to differentiate visual objects presented in ‘visual clutter’ [5,6]) and may be poorer at extracting signals from noise [5–7] as compared to the fovea. This means that visual processing remains poor, even after spatial resolution limitations are addressed with appropriate magnification. Therefore, the reduction of crowding could be an effective rehabilitation strategy. Perceptual learning studies in individuals with normal vision [8,9] and patients with central vision loss [10,11] have reported improvements in peripheral visual function with training. However, perceptual learning requires extensive training and gains are typically specific to the trained stimulus. Nevertheless, these findings indicate that the neural mechanisms contributing to crowding in peripheral vision remain modifiable.

Non-invasive brain stimulation techniques such as anodal direct current stimulation (a-tDCS) offer an alternative approach to improving peripheral visual function [12,13]. a-tDCS alters the excitability and neurochemical environment of brain regions close to the stimulation site. In particular, anodal tDCS (a-tDCS) increases cortical excitability and reduces the concentration of the inhibitory neurotransmitter GABA[14]. Using a-tDCS over the visual cortex, we have recently demonstrated that collinear inhibition can be reduced in observers with normal vision[15]. Collinear inhibition (also called lateral masking) refers to the impaired detectability of a visual target when it is flanked by similarly oriented high contrast targets and is one of the low-level mechanisms involved in visual crowding [16]. Building on this previous work, in this case series of three patients with CVL, we tested whether a-tDCS could be used to modulate the neural mechanisms that contribute to crowding in a clinical population. We observed that, compared to sham (placebo) stimulation, a-tDCS reduced collinear inhibition at the PRL in all three participants providing new proof of concept evidence for the use of a-tDCS to improve vision in patients with central vision loss.

## Materials and Methods

5 participants (4 females; age: 55±22 years) diagnosed with macular degeneration agreed to participate in this study. Participants were excluded if they had multiple PRLs and any contraindications to transcranial electrical stimulation (See Supplementary Material). Two participants were excluded due to a history of epilepsy and a preexisting heart condition. Data for the remaining three participants were collected at the School of Optometry and Vision Science, University of Waterloo, Waterloo, Canada (n = 2) and the Envision Research Institute, Wichita, USA (n =1). All participants provided written, informed consent. The study was approved by the University of Waterloo and Wichita State University research ethics committees. All the procedures involved in this research adhered to the tenets of the Declaration of Helsinki. All participants had a central scotoma which was determined using the Amsler Grid test and a microperimeter (OPKOS OCT/SLO) as shown in Figure-1.

**Figure 1:**
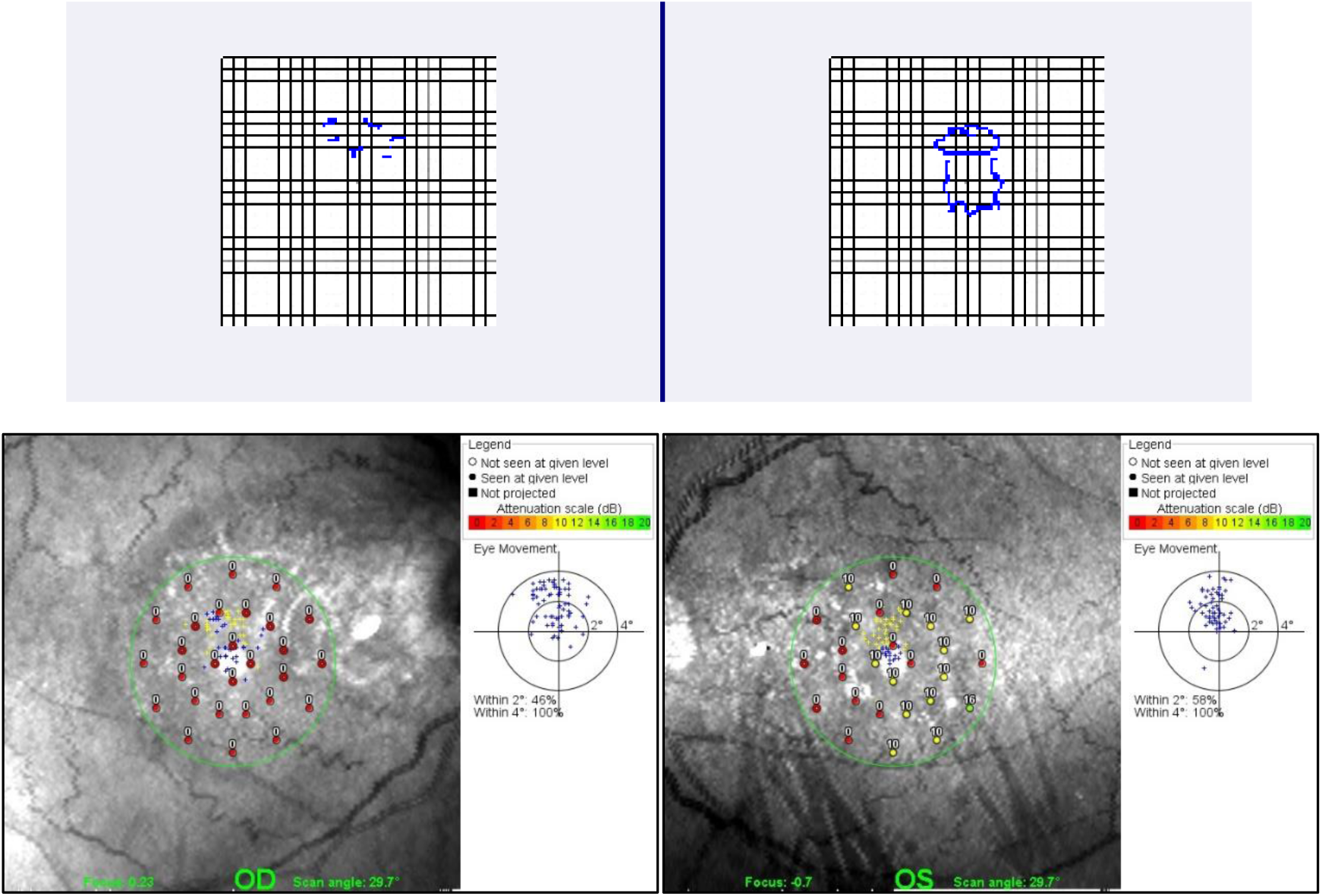
A sample Amsler grid test for S1 (top) and microperimetry image from S2 (bottom) showing a central scotoma.

A sham controlled, single-blind study design was used to test the effect of visual cortex a-tDCS on collinear inhibition in patients with central vision loss. Visual stimuli were presented on an LCD monitor that was placed at 100cm from a chin/forehead rest. Participants were instructed to fixate a central fixation cross (0.5°) and provide responses by pressing arrow keys on a keyboard. The target visual stimulus was a central Gabor patch (σ: 5, spatial frequency: 7 cpd) (Figure-2). Two flanker Gabor patches (σ: 5, spatial frequency: 7 cpd and a fixed contrast of 0.8) were placed above and below the target. Based on pilot observations, colinear inhibition was induced using target/flanker separations of 3λ for S1 and S3 and 2λ for S2 whose PRL was relatively closer to the fovea [10]. A larger target/flanker separation of 6λ was also tested as an additional control condition. Larger target flanker separations induce collinear facilitation, the opposite to collinear inhibition [10,15]. Participants were encouraged to adopt their habitual head position so that they could see the fixation cross clearly. A 2-alternative-forced-choice detection task was used to measure the contrast threshold for the central Gabor patch. The initial contrast of the central target was set at 0.5 and contrast was varied using a 2-down and 1-up staircase method with a fixed step size of 0.05. The staircase was terminated either after 60 trials or 5 reversals. The mean of the final four reversals was taken as the contrast threshold for the staircase. Each threshold measurement was the mean of two staircases. Throughout the experiment, fixation was monitored using an infrared video based eyetracker (EyeLink systems, SR Research). All participants received a practice period before the actual measurement of contrast thresholds using a fixed target contrast of 0.5. The practice measurements were repeated until the participant fully understood the task to minimize the effect of task learning.

**Figure 2:**
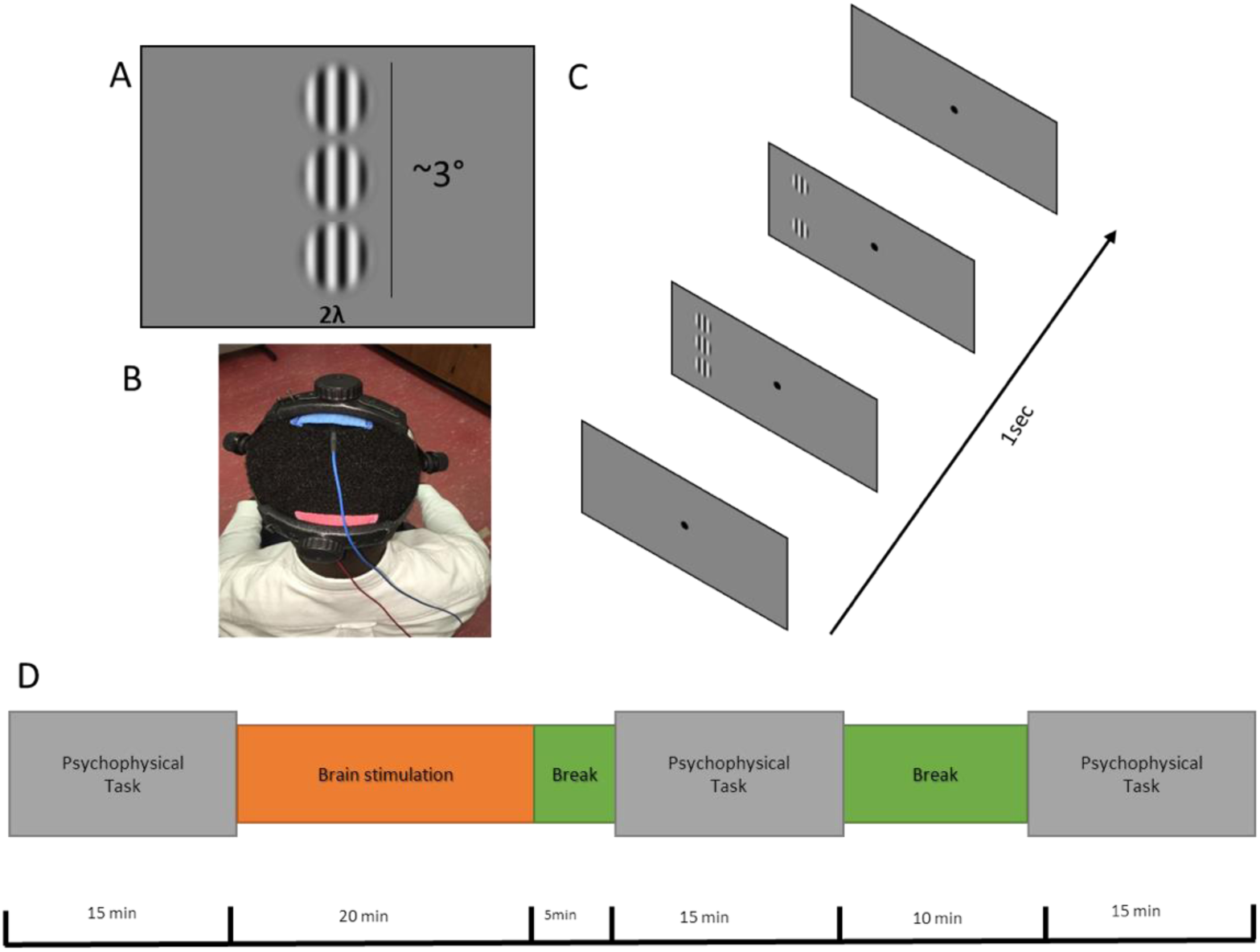
Visual stimuli and experimental design

tDCS was administered using a commercially available DC Stimulator (NeuroConn MC). The electrical current was administered using two 5cm x 5cm rubber electrodes placed inside a saline soaked sponges. The electrodes were secured using a head strap and placed over Oz (anodal electrode) and Cz (cathodal electrode) based on the 10-20 EEG electrode positioning system. Active stimulation involved 2mA of anodal tDCS for 20 minutes. Sham stimulation involved a 5 second ramp up of current immediately followed by a 5 second ramp down with the electrodes left in place for 20 minutes. Active and sham stimulation visits took place on two different days and each visit was separated by at least 48 hours. Participants were masked to the type of stimulation. The order of stimulation was also randomized. During each visit, all participants completed 4 sessions of contrast threshold measurements: pre-, during-, 5min post- and 30min post-stimulation (Figure-2). In addition, during each visit, participants were asked to complete adverse effects questionnaires before and after the stimulation sessions.

## Results

All participants completed the two study sessions without reporting any adverse effects. However, participant S3 could not complete the 6λ viewing condition. None of the participants were able to guess the stimulation type correctly. Table-1 shows the mean contrast threshold values and staircase reversal standard deviations for every participant for the collinear inhibition and facilitation viewing conditions. For each participant, the mean contrast threshold for the during-, 5min post- and 30min post-stimulation periods was normalized by subtracting the within session, baseline pre-stimulation contrast threshold. Figure 3 shows the mean normalized contrast thresholds for the collinear inhibition condition for the active and sham stimulation sessions. A reduction of collinear inhibition is apparent for the active but not the sham stimulation conditions for all participants. With a single session of a-tDCS, an average 56% (38% to 76%) reduction of collinear inhibition was sustained after 30min of stimulation.

**Table 1:** Mean contrast threshold values (db) and SD of reversals for each participant

| Subject ID | Inhibition |  |  |  |  |  |  |  |
| --- | --- | --- | --- | --- | --- | --- | --- | --- |
|  | Anodal |  |  |  | Sham |  |  |  |
|  | BL | DS | PS5 | PS30 | BL | DS | PS5 | PS30 |
| S1 | 0.470±0.06 | 0.270±0.08 | 0.245±0.05 | 0.113±0.05 | 0.350±0.10 | 0.450±0.01 | 0.360±0.05 | 0.270±0.01 |
| S2 | 0.373±0.04 | 0.383±0.07 | 0.228±0.06 | 0.230±0.05 | 0.408±0.06 | 0.408±0.10 | 0.475±0.07 | 0.375±0.05 |
| S3 | 0.433±0.06 | 0.144±0.05 | 0.244±0.07 | 0.196±0.11 | 0.235±0.06 | 0.306±0.08 | 0.206±0.05 | 0.195±0.06 |
|  | Facilitation |  |  |  |  |  |  |  |
|  | Anodal |  |  |  | Sham |  |  |  |
|  | BL | DS | PS5 | PS30 | BL | DS | PS5 | PS30 |
| S1 | 0.215±0.02 | 0.283±0.08 | 0.095±0.03 | 0.143±0.04 | 0.183±0.09 | 0.268±0.07 | 0.129±0.03 | 0.143±0.04 |
| S2 | 0.060±0.03 | 0.053±0.03 | 0.058±0.03 | 0.045±0.01 | 0.093±0.05 | 0.038±0.03 | 0.015±0.02 | 0.037±0.04 |

**Figure 3:**
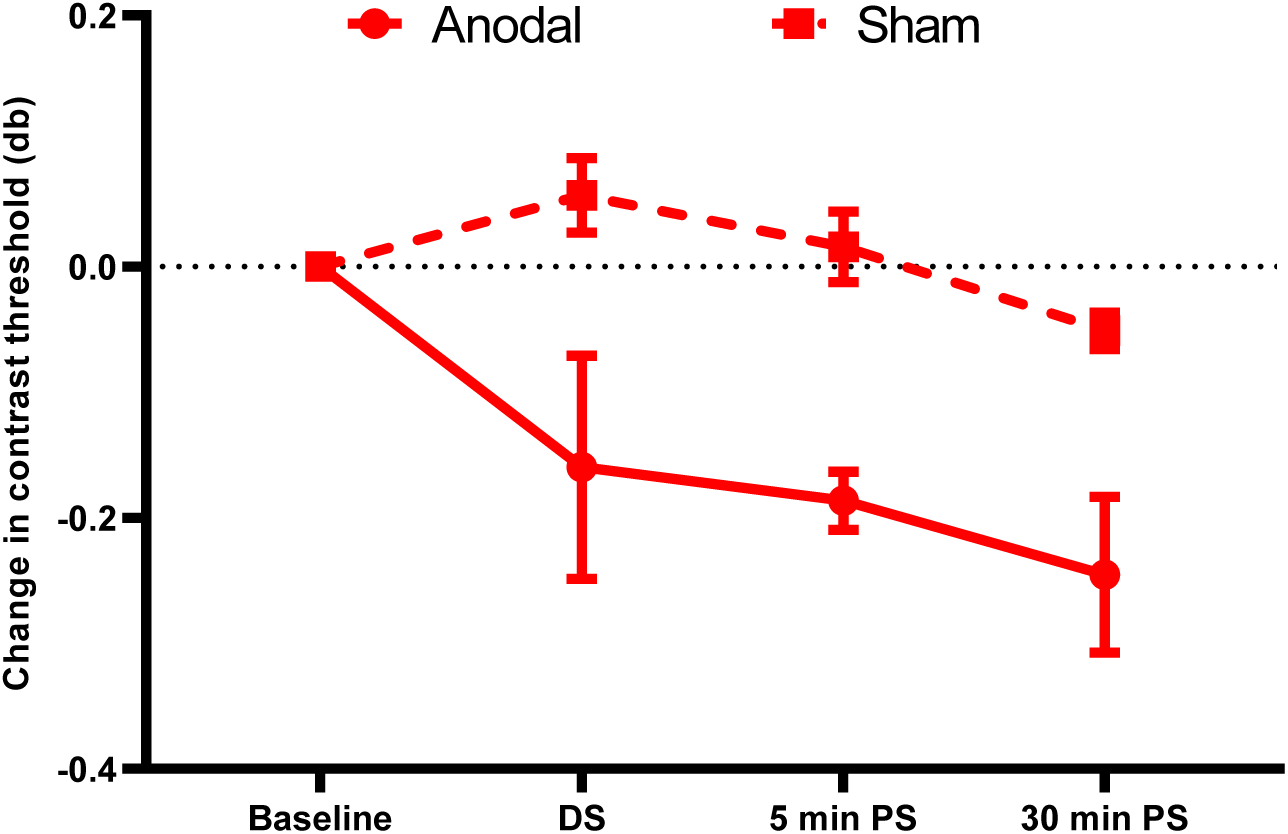
Change in contrast threshold for anodal and sham tDCS. The solid line and circled markers represent the active anodal stimulation whereas the dotted line and square markers represent the sham stimulation. Abbreviations: DS – during-stimulation; PS – post-stimulation.

## Discussion

Crowding is a primary cause of visual impairment at the PRL. Therefore, reduction of crowding can be an effective means for visual rehabilitation. Previous studies have used perceptual learning for this purpose. For instance, Chung (2007)[17] trained eight observers with central vision loss using a trigram paradigm and observed a 38% reduction in crowding and an 8% improvement in reading performance after about 6000 trials[5]. However, the effect of perceptual learning can be very specific, meaning that the improvements in vision do not necessarily transfer to tasks other than the training task. The extensive training required for perceptual learning might be difficult for patients, especially if improvements are mostly task-specific. Our preliminary case series results provide proof of concept that it is possible to alleviating crowding for patients with macular degeneration by directly altering cortical processing with non-invasive brain stimulation. This is consistent with our recent study demonstrating reduced collinear inhibition in observers with normal vision during and after applying a-tDCS to the visual cortex. Together, these observations provide a foundation for larger studies of the potential for a-tDCS to improve visual functioning in patients with CVL and provide a foundation for larger studies to employ a-tDCS either alone or in combination with other interventions such as perceptual learning or low-level electrical stimulation of ocular structures [18]. Recent technological advances improving the accessibility of a-tDCS can now be delivered as a home based therapy (for instance, https://soterixmedical.com/research/remote) meaning that such studies can be more easily accomplished.

## Acknowledgement

This research was funded by the LCI Foundation (RNR), CFI grant 34095(BT) and NSERC grants RPIN-05394 and RGPAS-477166 (BT).

